# Self-association of a nucleoid-binding protein increases with macromolecular crowding in *Escherichia coli*

**DOI:** 10.1101/2023.02.23.529735

**Authors:** Theodoros Pittas, Arnold J. Boersma

## Abstract

Many proteins self-associate to achieve function. Macromolecular crowding enhances protein self-assembly in buffer experiments with added crowders, and crowding could therefore regulate protein function and organization in cells. In eukaryotic cells, protein condensation has been shown to increase with crowding. However, it is unclear what the effect of crowding is on native protein self-assembly in the highly crowded *Escherichia coli* cell. To determine the role of crowding in the self-assembly of a native protein, we study here the nucleoid-binding H-NS in *E. coli* and alter macromolecular crowding using a set of perturbations. We followed H-NS self-assembly using a FRET-based method for determining intermolecular interactions with a single genetic intervention. In dilute cell lysate, we see that H-NS self-assembly increases with salts, macromolecular crowding, and its own concentration. In *E. coli*, the oligomerization increases with crowding. We see that the response of H-NS oligomerization to a sudden crowding change is not immediate but requires time to adapt. Our findings implicate that in-cell crowding affects intracellular organization by promoting self-assembly.

## Introduction

Self-assembly or oligomerization can endow a protein with a function, such as a nucleolus for ribosome synthesis or the cytoskeleton for cell shape and organization. In contrast, others have less clear functional relevance and may be pathogenic, such as α-synuclein aggregates in Parkinson’s disease.^1^ Self-assembly is governed by the architecture of the interacting surfaces of the proteins, which is, in turn, mediated by the local physicochemical conditions such as ionic strength, pH, temperature, and macromolecular crowding. In the *E. coli* cell, the concentration of macromolecules ranges between 200-300 mg/mL,^2,3^ and this high crowding should be relevant to any self-assembling protein. Theory and experiments in buffer showed that oligomerization and aggregation could be enhanced by the volume crowders take up.^4–6^ In principle, as crowders are depleted from the smaller space between two proteins, the generated pressure difference pushes the two proteins together. This effect, known as the depletion force, depends on various parameters, including the crowder number density, the crowder shape and volume, and the shape and volume of the depleted zone between the two interacting proteins. Moreover, any associative or repulsive interaction between the crowder and the self-assembling proteins will modulate the effect of the crowders. Therefore, while oligomerization is clearly important and should be enhanced by macromolecular crowding, the inherent complexity of the in-cell crowding effect does not allow for predicting its consequences on protein self-assembly in the heterogeneous intracellular environment.

The effect of crowding on proteins in living cells can be extracted from perturbations of the macromolecular crowding through stress conditions. For example, mechanical stress, osmotic stress, or energy depletion may alter the crowding,^7–9^ of which hyperosmotic stress is most often used, albeit this increases the concentration of all components, including the protein under investigation. Recent evidence on the effect of macromolecular crowding on protein self-assembly in cells was provided for condensate formation in eukaryotic cells: Condensates of ASK3 were induced by macromolecular crowding under hyperosmotic stress,^10^ while increasing ribosomal crowding enhanced phase separation of a model condensate,^11^ and WNK kinases formed condensates upon injecting Ficoll into the cells.^12^ As the *E. coli* cytoplasm has a higher biomacromolecule content, one would expect higher crowding effects in such cells. Although experiments with purified protein in a buffer often require the addition of crowders to mimic the crowding in *E. coli*,^13–15^ the role of the crowding inside *E. coli* on the self-assembly of its proteins has not been determined.

A well-studied protein that functionally self-associates is the Histone-like Nucleoid Structuring protein (H-NS) in *E. coli*. This protein is highly abundant, with >20,000 copies per cell.^16^ H-NS is a Nucleoid-Associating Protein (NAP) involved in DNA organization and topology.^17,18^ It binds and oligomerizes along DNA, forming bridges, loops, or lateral filaments.^19–23^ H-NS oligomerizes at intrinsically-curved sequences or AT-rich regions.^18,24–26^ The formation of such complexes is linked with supercoiling suppression and transcription regulation.^18^ H-NS consists of an N-terminal oligomerization domain, a C-terminal DNA-binding domain, and a disordered linker region in between. The self-assembling behavior of this protein has been studied in detail in buffer and cells. In buffer solutions, the N-terminus and specifically the amino acids 1-64 forms lower-order oligomers, such as dimers or trimers.^24,27–30^ Amino acids 65-89, on the other hand, are required for higher-order oligomers with less defined stoichiometry.^18,27,29,31–33^ The DNA-binding domain does not participate in oligomerization.^24,27,28^ The degree of H-NS oligomerization is concentration-dependent in vitro. Next to this, ions, such as Na^+^, K^+^, and Mg^2+^, affect H-NS self-association and the conformation of the H-NS-DNA complexes.^19,20^ Magnesium and other factors can promote an open H-NS conformation, which is DNA bridging competent, while the closed conformation is not. In this manner, it may regulate transcription. In general, an increased ion concentration affects the conformation of members of the bacterial histone family, rendering them more extended and flexible.^34^ Despite these well-described sensitivities of H-NS to various solution conditions, the sensitivity of its oligomerization to in-cell macromolecular crowding is not known.

Here, we show that H-NS assembly is sensitive to the macromolecular crowding inside living *E. coli* cells by perturbing crowding with recently developed methods.^35^ We obtain insight into the protein self-assembly behavior of H-NS using intermolecular FRET.^36^ We see that the macromolecular crowding enhances the oligomerization of H-NS in both dilute lysate and in *E. coli*, although the H-NS oligomerization in *E. coli* requires time to respond to a sudden crowding change.

## Results

### In vitro H-NS self-association is sensitive to salts, crowding, and its own concentration

To follow the self-assembly or oligomerization of H-NS, we used a genetically-encoded FRET pair on a single genetic construct optimized for intermolecular FRET.^37^ We recently showed that this method allows straightforward determination of protein self-organization in a precise and high-throughput manner.^36^ The method relies on a tight fusion between an mVenus and mCherry (VC) with low intramolecular FRET due to the restricted motion between the two fluorophores. In contrast, the formation of condensates, aggregates, and oligomers should increase FRET efficiency through intermolecular FRET.^36,38^ The readout depends on the intermolecular distance between the FRET pairs fused to the target proteins, which in turn is a function of the density and structure in the assembly.^38,39^

We fused H-NS to the VC domain via a short 5-Glycine linker (Figure 1). The glycine linker prevents hindrance between the VC and H-NS, so that they both fold and function properly. We fused the C-terminus of the H-NS protein to mVenus. To capture the behavior of the individual domains, we created two mutants of the oligomerization domain based on the literature.^31^ We selected the mutants containing the amino acids 1-64 and 1-86, which we named H-NS_64_-VC and H-NS_86_-VC. H-NS_64_ has been shown to form dimers or trimers in buffer. H-NS_86_, on the other hand, forms dimers of dimers and higher-order oligomers.

**Figure 1.**
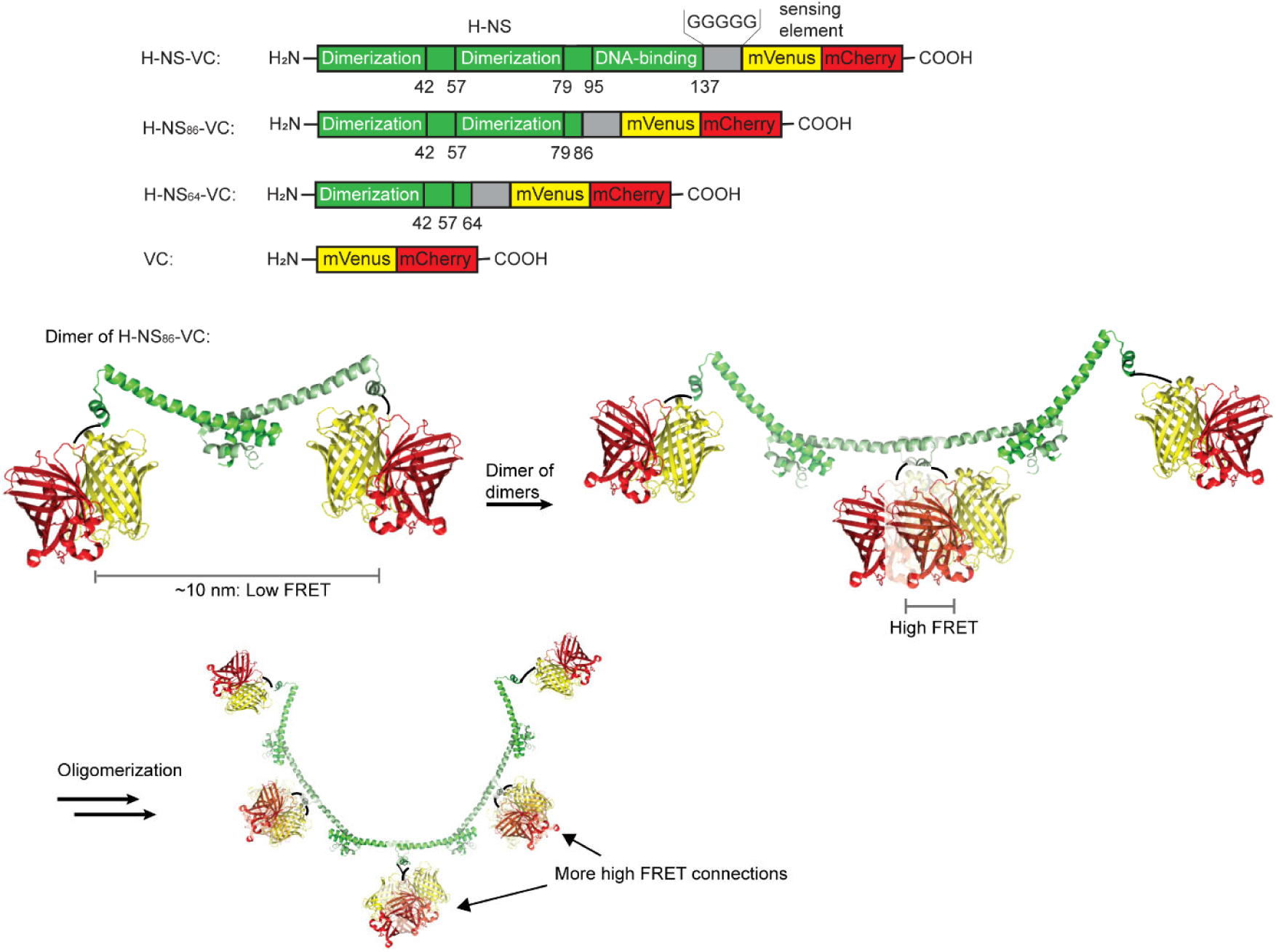
Design and working model of H-NS oligomerization sensing with an intermolecular FRET module. An mVenus/mCherry (VC) domain that displays low intramolecular FRET is fused to wt-H-NS, H-NS_1-86_, and H-NS_1-64_ to capture the role of DNA binding, oligomerization, and dimerization. The monomers dimerize at their N-termini, inducing a low intermolecular FRET between the VC domains at the C-termini. Dimerization of the dimers at these C-termini induces high FRET between the VC domains. Further oligomerization increases the number of high FRET connections relative to the low FRET connections.

Using this methodology, we can study H-NS oligomerization with FRET. It was shown that upon dimerization at its N-termini, the radius of gyration of a truncated (amino acids 1-83) in the *Salmonella typhimurium* H-NS dimer is 3.9 nm, as determined from SAXS.^33^ We determined the distance between the C-termini, where the fluorophores are attached, as ∼10 nm from its crystal structure (PDB: 3NR7) with Pymol. These values are beyond the range of the Förster distance of ∼5 nm for fluorescent proteins, at which there is 50% FRET efficiency, and we, therefore, expect a low FRET for the dimers. However, when the dimers dimerize at their C-termini to provide the proposed linear structures,^33^ the fluorescent proteins are in very close proximity and give maximal FRET. Moreover, if oligomerization continues, the number of high-FRET contacts will increase while there will always be two low FRET N-termini remaining, therefore further increasing the FRET efficiency. Hence, an increase in the oligomerization state translates into an increase in FRET efficiency.

To characterize H-NS self-association in vitro, we overexpressed the gene encoding the protein from the pBAD plasmid in *E. coli* BL21(DE3) and lysed the cells so that we could controllably perturb the H-NS. We determined the fluorescence of the constructs by scanning confocal fluorescence microscopy without further purification in dilute cell lysate. We treated the lysate with DNAse-I to homogenize the solutions. However, DNA fragments may remain and take part in the assembly in the case of H-NS-VC. Indeed, we see a fluorescent signal in the loading well with in-gel fluorescence of lysates run on native PAGE gels that may correspond to H-NS-VC/DNA complexes (Figure S1). Using these dilute lysates, we studied conditions relevant to oligomerization, such as macromolecular crowding (15% w/w Ficoll-70) and increased salts (50 mM magnesium chloride or 500 mM potassium glutamate) on H-NS-VC and the H-NS_86_-VC mutant. Salts are known to influence H-NS self-association and DNA-binding behavior.^19^ In addition, we tested the concentration dependence of the constructs.

When analyzing the microscopy images of the H-NS-VC and H-NS_86_-VC in the dilute cell lysates, we see clearly visible foci that reach up to 1.5 μm in diameter (Figure 2a,b), with a significant fraction in the dilute phase. As a measure of FRET efficiency, we excited the donor (mVenus) and divided the resulting acceptor (mCherry) emission by that of the donor. This FRET ratio is similar for H-NS-VC and H-NS_86_-VC foci, 0.07±0.002 and 0.073±0.004, respectively, showing that the DNA binding domain does not change the oligomerization behavior. Moreover, the FRET ratios in the foci are similar to the solution outside the foci: 0.082±0.007 and 0.068±0.03 for H-NS-VC and H-NS_86_-VC, suggesting that the proteins are in a similar oligomerization state. We see a large increase in the FRET ratio in both the foci and dilute phase, increasing from 0.07±0.002 to 0.16±0.03 (±s.d.,n=3) and from 0.08±0.005 to 0.22±0.06, respectively, within 30 min after the addition of the crowder, the salts, or after an increase in protein’s own concentration, for both constructs (Figure 2 c-d, f-g). The ratios remained stable for at least 3 h. Hence, the self-assembly structure of these proteins changes to the same extent for all additives or with protein concentration in a switch-like manner. Given the design of the sensor (Figure 1), we hypothesize that this high increase in FRET represents the formation of oligomers from the dimers.

**Figure 2.**
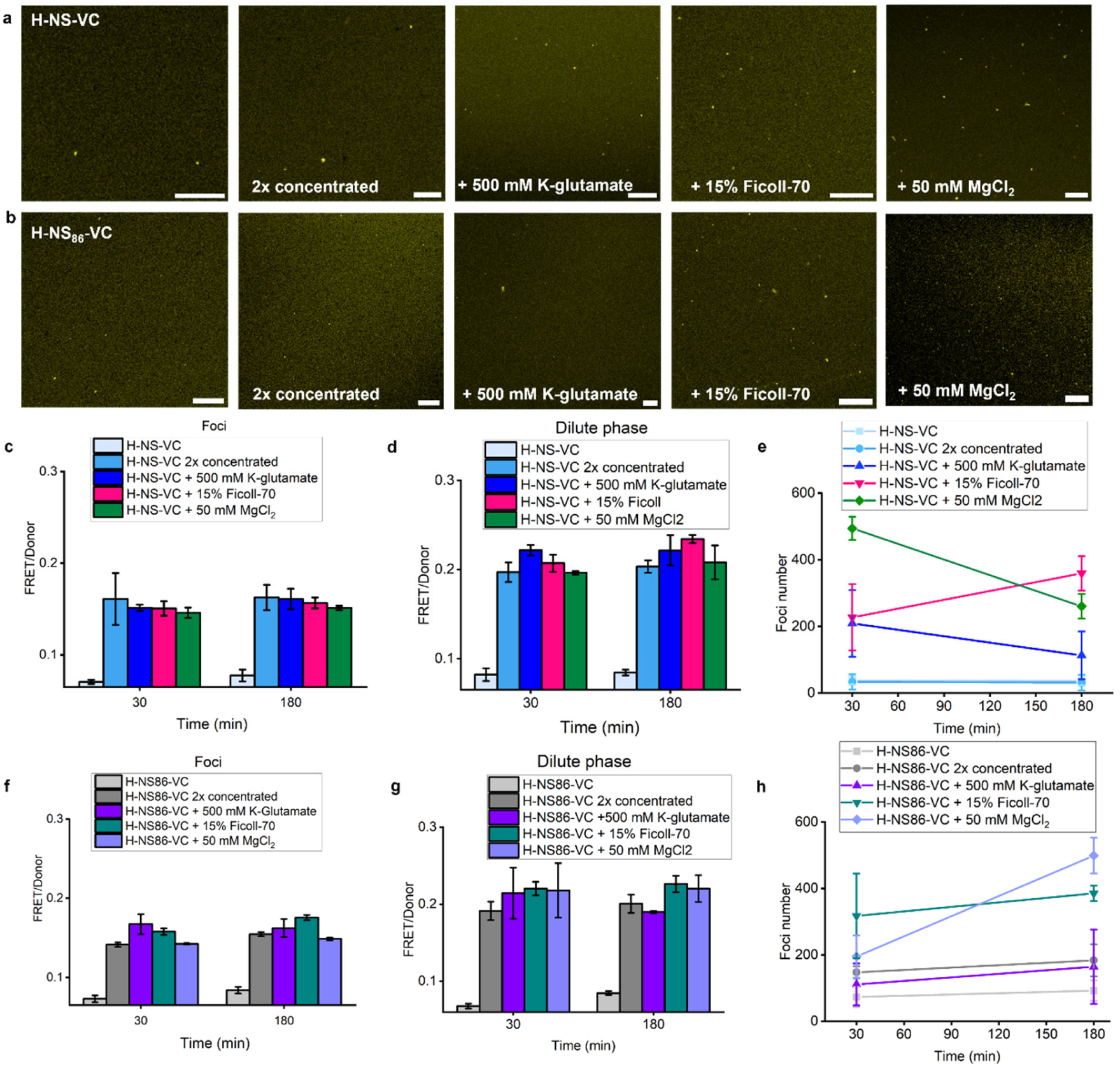
Self-association of VC constructs increases in dilute lysates under conditions of increased crowding, ionic concentration, or protein concentration. Representative fluorescence confocal images of the Donor channel of (a) H-NS-VC and (b) H-NS_86_-VC after increasing the protein’s own concentration or adding 15% w/w Ficoll-70, 500 mM potassium glutamate, or 50 mM magnesium chloride. FRET/Donor ratio in (c) H-NS-VC foci, (d) H-NS-VC dilute phase, (f) H-NS_86_-VC foci, and (g) H-NS_86_-VC dilute phase. (e) H-NS-VC foci number (h) H-NS_86_-VC foci number. The data is the average over three independent experiments, and the error bars are the standard deviation. The size of the scale bar is 15 μm.

When analyzing the macroscopic foci properties, we see that without additives, the H-NS-VC forms two times fewer foci than H-NS_86_-VC (Figure 2e,h), and the dilute phase of H-NS-VC appears slightly denser than H-NS_86_-VC. For H-NS-VC, the foci number increases with all additives but not when increasing its own concentration, while H-NS_86_-VC forms more foci in all cases. In contrast to the relatively uniform behavior in foci numbers, their sizes were highly additive-specific (Figure 3). In the case of the crowding agent Ficoll-70, the increase in foci number is associated with smaller foci for both constructs. This state was not stable and slowly redistributed towards larger foci. Hence, the macroscopic properties of the foci do not reflect the oligomerization structures inferred from FRET, which are more binary.

**Figure 3.**
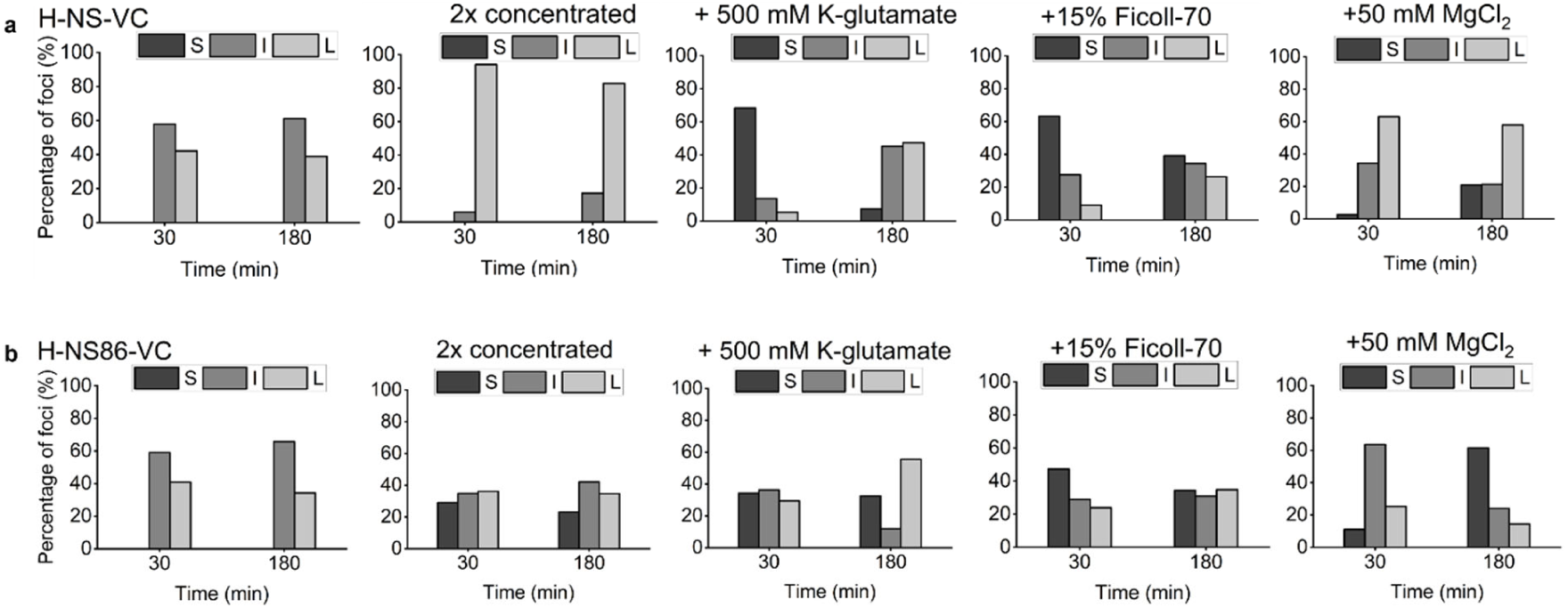
VC-constructs foci size varies in vitro under conditions of increased protein concentration, crowding, and ionic strength. Shown here are the histograms of the foci size distribution, classified as small (S), intermediate (I), and large (L) for (a) H-NS-VC and (b) H-NS_86_-VC. Each panel contains all foci from three biological replicates for each construct and each condition. Small foci size is <0.1 μm, intermediate 0.1-0.2 μm, and large >0.2 μm.

### H-NS self-assembly in *E. coli* follows crowding

To be able to determine if H-NS oligomerization increases with crowding in its native crowded context, *E. coli*, we studied its behavior first in exponentially growing cells. To this end, we expressed the constructs in *E. coli* BL21(DE3) using an arabinose inducible promoter to control the expression levels. After inducing for 3 h with 5 μM arabinose while maintaining the cells in the exponential growth phase (OD_600_ = 0.1-0.3), we imaged the cells by fluorescence confocal microscopy. The constructs did not show the same spatial distribution (Figure 4a): H-NS-VC contains the C-terminal DNA-binding domain and is mostly where the nucleoid can be expected and is consequently excluded from the poles.^31,40^ Hence, VC-fusion does not inhibit its DNA-binding capability. The H-NS_64_-VC, which has been shown to form dimers and trimers in buffer,^33^ is distributed evenly in the cytoplasm, similar to the VC control. In contrast, H-NS_86_-VC, which was shown to form higher oligomers in buffer, formed a single focus per cell, which is excluded from the nucleoid region and resides in a cell pole. Each construct gave these distribution patterns consistently in each cell. Hence, the localization of H-NS and its truncated mutants reflects reported observations in buffer despite the fusion to the VC domain.

**Figure 4.**
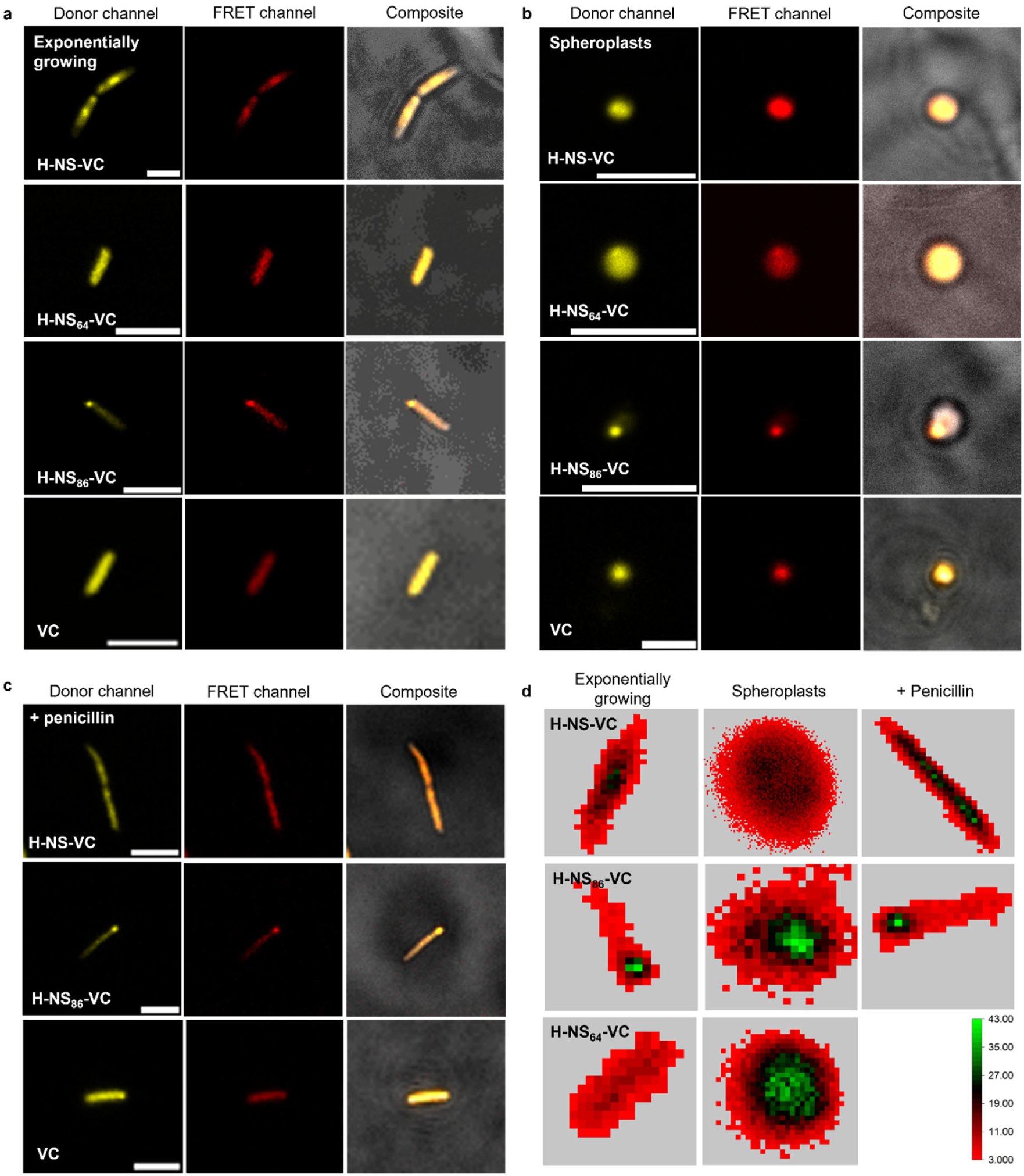
Representative fluorescence confocal images of the Donor and FRET channels and of the brightfield channel overlayed with the former channels to show the localization of the VC constructs in *E. coli*. (a) The exponentially growing cells, (b) lysozyme-spheroplasts, and (c) penicillin G-treated cells (1 min after treatment with 0.5 mg/mL Penicillin G) after expression of H-NS-VC, H-NS_64_-VC, H-NS_86_-VC, and VC. (d) Heat maps of the FRET channel of a representative exponentially growing cell, lysozyme-spheroplast, and penicillin-treated cell (1 min after treatment with 0.5 mg/mL Penicillin G) of the H-NS-VC, H-NS_64_-VC, and H-NS_86_-VC normalized to the background, which is the fluorescence intensity values of media used for each condition (on average 2.5). The color bar denotes fluorescence intensity. The scale bars are 4 μm.

The FRET ratios of the constructs are relatively similar, with the H-NS_64_-VC and H-NS_86_-VC giving slightly higher ratios (0.126±0.006 and 0.132±0.007, respectively) than H-NS-VC (0.105±0.005) (Figure 5a). Hence, even though H-NS_64_-VC and H-NS_86_-VC have distinct spatial localization patterns, the structures are similar at the 2-10 nm FRET distance range. Indeed, when comparing the ratios in the foci and the cytoplasm for H-NS_86_-VC (Figure 5a), we see similar FRET ratios, which reiterates the independence of the oligomerization behavior on the localization. Nonetheless, the lower ratio of H-NS-VC suggests that the DNA binding domain affects the H-NS oligomer structure in an intracellular environment but not in dilute lysates, where all the constructs provide similar FRET. To verify the ability to determine the oligomerization state, we subjected the cells to a 6 min heat shock at 42 °C. It was previously shown that temperature increases the oligomerization state of H-NS,^41^ and here we see an increase in the H-NS-VC ratio to 0.162±0.003 (Figure S2) indeed. The heat shock does not change the localization of the H-NS, and therefore DNA dissociation does not cause the ratiometric change. Together, the FRET ratios of the different constructs are similar in *E. coli* irrespective of the location but decrease somewhat due to the DNA-binding domain.

**Figure 5.**
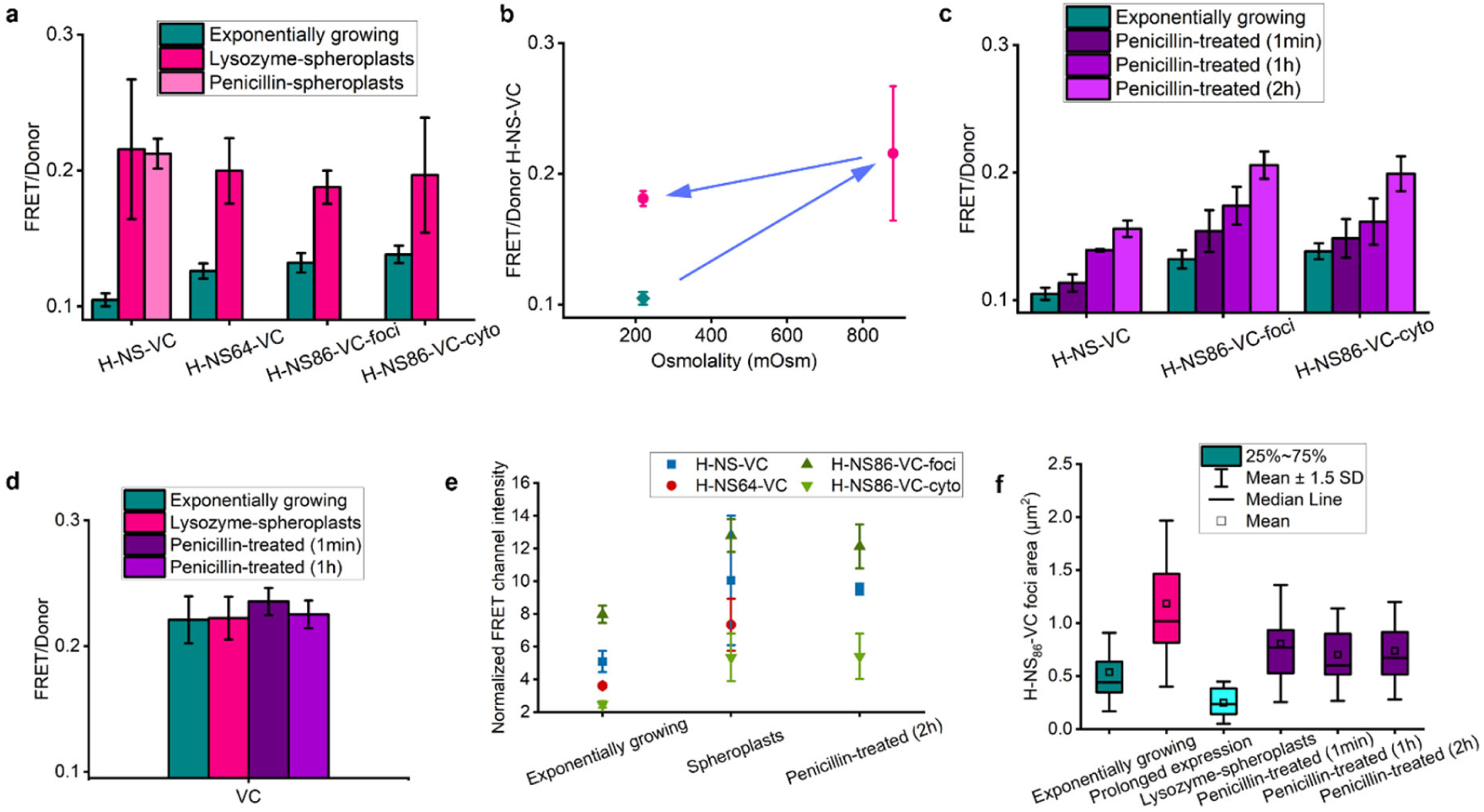
Spheroplast formation and penicillin treatment increase the FRET/Donor of the VC constructs as analyzed from fluorescence confocal microscopy. (a) Increase in FRET/Donor for the VC constructs in spheroplasts, in comparison with exponentially growing cells. (b) The effect of medium osmolality on H-NS self-association in spheroplasts. We decreased the external osmolality of spheroplasts (880 mOsm) to 220 mOsm, which is equal to exponentially growing cells in MOPS medium. Blue arrows are the order of treatment. Green = exponentially growing cells, pink = spheroplasts. (c) Increase in FRET/Donor of the VC constructs after treatment with 0.5 mg/mL Penicillin G followed in time. (d) FRET/Donor ratios of the VC control after the perturbations. (e) FRET channel intensity normalized to the average background signal for each construct for exponentially growing cells, lysozyme-spheroplasts, and penicillin-treated cells (∼ 90 cells per condition and construct). (f) Box plot of the H-NS_86_-VC foci surface area for the different conditions (∼ 100 cells per condition). All data is the average of three independent biological replicates. Error bars are the standard deviation.

Next, we determined the effect of crowding on the H-NS oligomerization. We used recently identified conditions that drastically increase crowding effects by forming spheroplasts of the *E*.*coli* cells using lysozyme/EDTA- and penicillin-based protocols.^35^ We see that after spheroplasting, the H-NS_86_-VC again forms foci, and the H-NS_64_-VC is again homogeneously spread throughout the cell (Figure 4b), while the H-NS-VC now occupies the entire cell as the nucleoid expands.^35^ All three constructs provide an increase in FRET ratio to roughly the same value for both spheroplasting protocols (Figure 5a). In contrast, the FRET ratio of the VC control did not change significantly (Figure 5d), excluding fluorescent protein artifacts. Hence, spheroplast formation, which drastically increases the macromolecular crowding, also increases the self-assembly of H-NS and its truncated mutants.

To determine if the enhanced self-assembly in spheroplasts is not due to an increase in ion concentration, we returned the spheroplast medium from 880 mOsm to a growth medium of 220 mOsm (Figure 5b), which should decrease the intracellular ion content to maintain isosmotic conditions with the environment. Under these conditions, the H-NS-VC ratio remains higher than exponentially growing cells (0.181±0.006, n=3), similarly as we previously found for macromolecular crowding.^35^ The retention of the higher H-NS-VC ratio implies that the ion concentration has little effect on the oligomerization.

Next, we determined if the change in volume and shape of the spheroplasts is responsible for a change in oligomerization: a reduction in volume increases the concentration of all components, including H-NS and molecules that interact with H-NS. Therefore, we added Penicillin G to exponentially growing cells (Figure 4c). We previously showed that penicillin treatment increased cell volume but also macromolecular crowding effects, likely due to a change in cellular organization.^35^ While the crowding increases immediately after penicillin addition, we see that the H-NS constructs show only a minor increase in their FRET ratios after 1 minute. However, the FRET ratio for both the H-NS-VC and the H-NS_86_-VC continues to increase for at least 2h incubation with Penicillin G (Figure 5c). This time dependence is not due to fluorescent protein maturation because the FRET for the VC control does not change. The oligomerization increases during the death phase and when the population doubles (Figure S3), showing little dependence on the growth phase. We can thus conclude that the reorganization of the H-NS to form higher oligomers lags behind the macromolecular crowding but eventually oligomerizes as expected from the crowding increase. Moreover, the oligomerization state of the H-NS does not follow the cell volume or shape.

Next, we assessed if the H-NS concentration determines the degree of oligomerization. The fluorescence in the FRET channel is the highest in the H-NS_86_-VC foci, followed by H-NS-VC, H-NS_64_-VC, and H-NS_86_-VC in the cytoplasm has the lowest intensity (Figure 4d, 5e) both before and after the perturbations. Hence, the protein concentration does not determine the FRET because this intensity difference is not reflected in the corresponding FRET ratios. Moreover, reducing the expression by lowering the arabinose content from 5 µM to 1 and 2 µM did not change the FRET ratios. We next analyzed if the FRET ratio depended on the intensity of single cells within a population. We took the sum of the donor and FRET channel as the best approximate for concentration because the small size of *E. coli* prevented accurate measurement with two laser excitations while keeping the cells in the same focus. When analyzing these single cells for all the perturbations, we see a variety of weak correlations, but there is no reliable correlation between the two parameters (Figure 6). Upon increasing the expression time from 3 h to 6 h, while maintaining the OD_600_ between 0.1-0.3, we see an increase in the FRET for all constructs, while the H-NS_64_-VC increases somewhat less (Figure S4). However, we see that the fluorescence intensity does not increase significantly (Figure 6), likely due to continued cell growth compensating protein synthesis, and, therefore, the H-NS-VC concentration cannot account for the increase in FRET with expression time. Hence, the H-NS concentration has, in comparison to crowding, little effect on the oligomerization.

**Figure 6.**
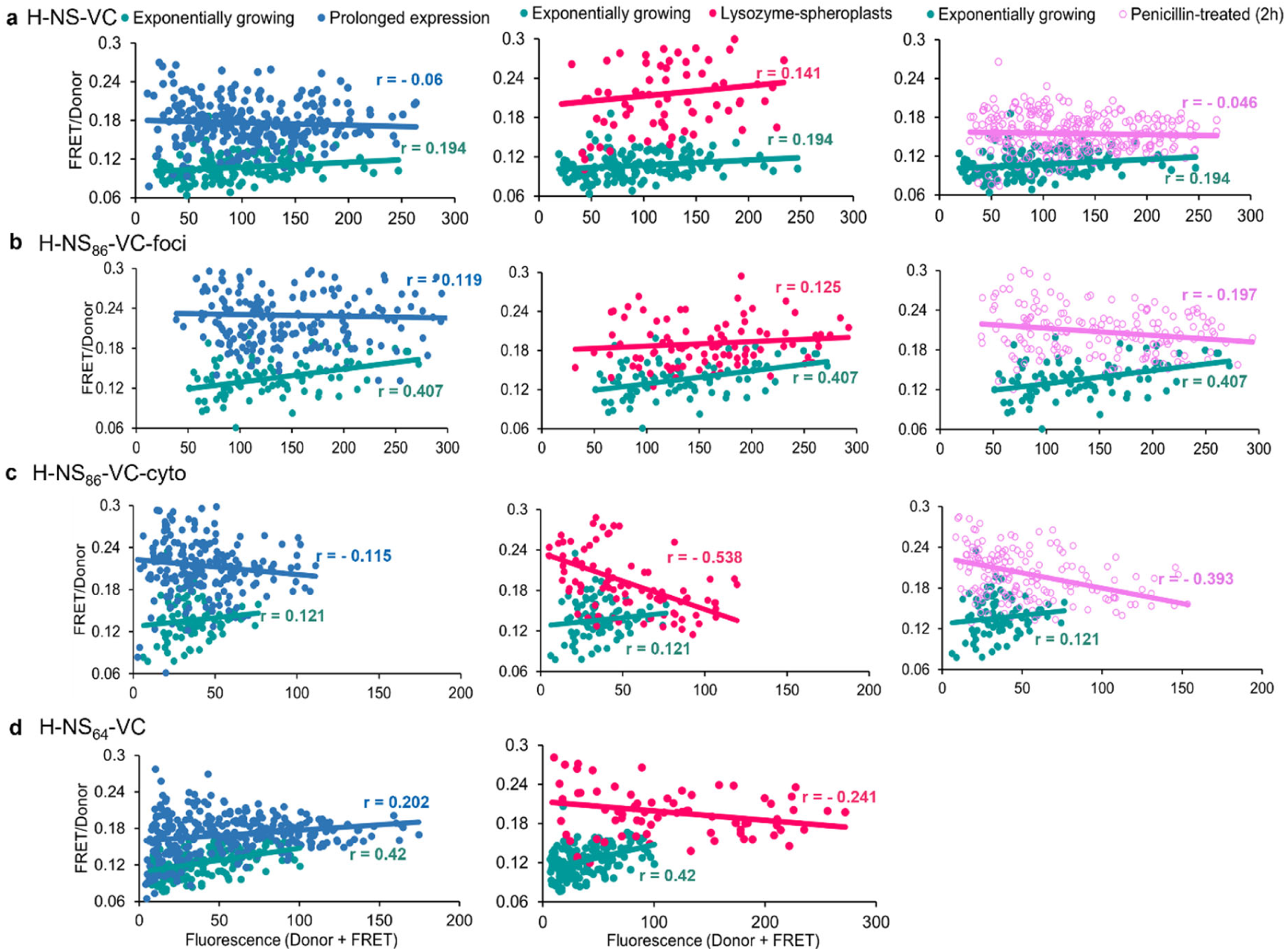
Self-association of the VC-fused constructs correlates poorly with protein concentration. Scatter plots of the FRET/Donor versus the fluorescence of the donor + FRET for single cells expressing (a) H-NS-VC, (b) H-NS_64_-VC, (c) H-NS_86_-VC foci and (d) H-NS_86_-VC cytoplasm. r = Pearson’s coefficient.

During the perturbations, we see a wide variety of H-NS_86_-VC foci diameters from which we cannot discern a clear trend. The foci size relates poorly to the H-NS_86_-VC concentration because the concentration of H-NS_86_-VC is higher in spheroplasts and penicillin-treated cells (Figure 5e), but the foci decrease in size for the former and do not change for the latter (5f). In contrast, the foci area of the H-NS_86_-VC foci after overexpression increases 2-3 times (Figure 5f). Thus, as in dilute lysates, the in-cell H-NS_86_-VC foci have a strong condition-dependent size distribution without a clearly discernible trend.

It is clear that the described conditions that induce high macromolecular crowding also enhance H-NS oligomerization. To determine if the degree of oligomerization and the amount of crowding positively correlate when compiling the different perturbations, we plotted the FRET ratios obtained from the H-NS probe oligomerization versus those previously obtained with a macromolecular crowding sensor (Figure 7).^35^ We included the data from exponentially growing cells, spheroplast formation with two different methods, an osmotic downshift of spheroplasts, and Penicillin G treatment followed in time. Next, we fitted the data versus a linear regression without the 1-minute Penicillin G data due to the time dependence of Penicillin G. We see reasonably good positive correlations between the two parameters ranging between r = 0.54-0.96 for the different constructs, especially when taking the different cell physiologies and sensitivity of H-NS to other parameters such as ionic strength and DNA binding, and time dependencies, into account. Moreover, we can clearly see that after 1 and 2 h with Penicillin G, there is an increase in H-NS oligomerization toward the linear fit expected from intracellular crowding. The correlation for H-NS-VC is less good than for H-NS_86_-VC and may require more time for the oligomerization state to stabilize from its lower FRET ratios in exponentially growing cells. In conclusion, despite the varying cell physiologies studied here, there is a positive correlation between the H-NS oligomerization state and the macromolecular crowding in *E. coli*.

**Figure 7.**
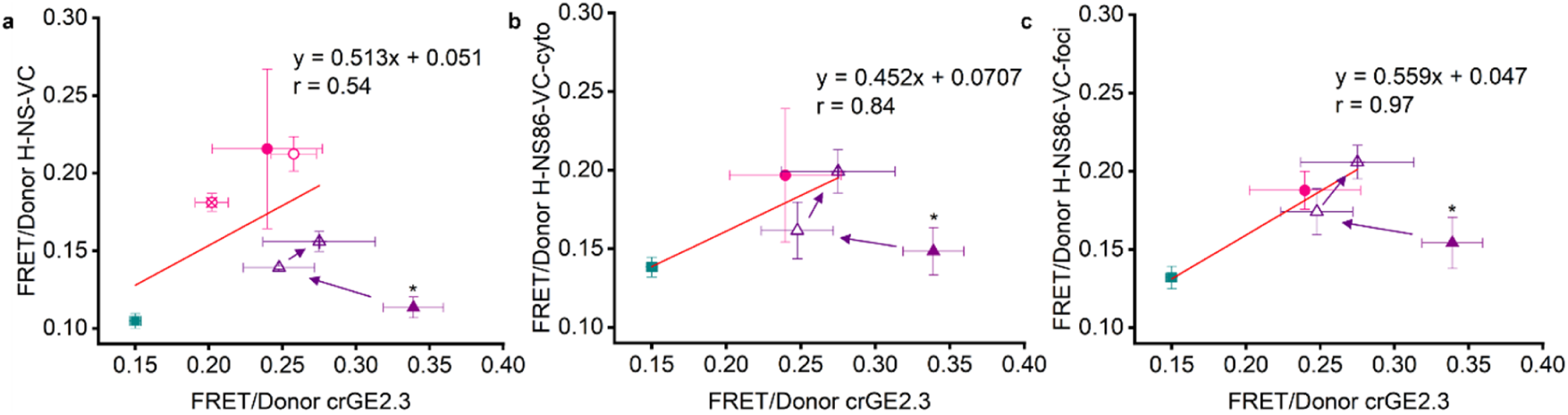
Self-association of H-NS-VC and H-NS_86_-VC correlates with macromolecular crowding. The FRET/Donor of (a) H-NS-VC, (b) H-NS_86_-VC cytoplasm, (c) H-NS_86_-VC foci, plotted versus the FRET/Donor of the crGE2.3 crowding sensor taken from Pittas et al.^35^ The data, without penicillin-treated cells at t =1 min (marked with an asterisk), was fit versus a linear regression (red line). The exponentially growing cells are represented by a green square, lysozyme-spheroplasts by a solid pink circle, penicillin-spheroplasts by a pink open circle, osmotically downshifted lysozyme-spheroplasts by a pink open circle with a cross, penicillin-treated cells at t = 1 min by a solid purple triangle, penicillin-treated cells at t = 1h by a purple open triangle, penicillin-treated cells at t = 2h by a purple open triangle with a cross. r = Pearson’s coefficient. Arrows indicate the time-dependent ratio of the VC constructs with Penicillin G. All data is the average of three independent biological replicates. Error bars are the standard deviation of three biological replicates.

## Discussion

In this study, we fused a domain for intermolecular FRET to the nucleoid binding protein H-NS to determine how its self-assembly depends on macromolecular crowding in its native host cell. We first show that H-NS self-assembly increases in dilute cell lysates after adding salts, Ficoll-70, or increasing its concentration. Inside cells, the localization depends on the H-NS mutant, while the H-NS oligomerization state is mostly uniform, with only a minor dependence of the oligomerization state on DNA binding. In *E. coli*, the H-NS self-assembly behavior was predominantly determined by the level of crowding. However, the H-NS self-assembly response was delayed behind the sudden crowding change.

We, and Ma *et al*., previously validated the VC domain for determining self-assembly with intermolecular FRET in mammalian and yeast cells.^36,37^ The major advantage of this method is that only a single genetic construct is needed. Here, we see that this approach also functions in *E. coli* because the FRET of the VC domain without H-NS remains unchanged under these perturbations. Therefore, the FRET changes are not due to fluorescent protein artifacts such as maturation but can be ascribed to oligomerization. Moreover, we see the expected oligomerization increase with temperature.^41^ In cells, we see less binary behavior of the oligomerization state as in vitro, although this may be due to the time dependence. As the constructs gave similar FRET ratios, conformational changes in H-NS, such as the open and closed conformations with its DNA-binding domain, likely do not play a major role. Co-oligomerization with native H-NS or other native proteins does not dominate the FRET, as the FRET was relatively independent of H-NS-VC concentration. We further see that its oligomerization behavior is modulated by its DNA-binding characteristics due to the lower FRET in the presence of the DNA-binding domain. While the VC domain consists of two bulky fluorescent proteins, it does not inhibit H-NS localization at the nucleoid where it is expected to localize, showing that the H-NS retains at least part of its functionality.^31,40,42^ Hence, the VC probing method is valid in *E. coli* and reports on H-NS self-assembly.

We see H-NS foci around the diffraction limit of ∼0.2 µm in dilute lysates and in cells for H-NS_86_-VC. As we measure at the diffraction limit, there are most likely also smaller clusters that we cannot image with a regular confocal microscopy setup. H-NS is known to form clusters or dense foci at the nucleoid,^31,40,42^ but those are larger and have a different shape than the spherical H-NS_86_-VC foci in cells or the H-NS_86_-VC and H-NS-VC foci in dilute lysates that we see. In the case of H-NS-VC in vivo, nucleoid binding will prevent the formation of the foci, while DNA digestion in dilute lysates likely allows the small foci to form. The size of the foci depends on the conditions in dilute lysates and in cells. For example, magnesium decreases the foci size of H-NS_86_-VC and H-NS-VC in dilute lysates: magnesium is known to bind to the glutamate residues of the H-NS oligomerization domain and prevents its closed conformation,^19,20,43^ providing a possible mechanism to alter foci properties. This may happen if magnesium leaks into the spheroplasts during their formation, albeit that many other mechanisms would allow explaining the smaller foci. However, throughout our data, the variation in foci size has little in common with the FRET values, possibly because of the different length scales where the clusters are seen and FRET distances, the latter of which is in the 2-10 nm domain. Hence, H-NS has a tendency to form clusters, but their properties are variable and do not represent the oligomerization state.

In vivo, protein oligomerization followed crowding. Macromolecular crowding has been previously shown to enhance protein oligomerization in buffer solutions: Linear oligomers formed by FtsZ became longer and more prevalent by the addition of 15% w/w protein crowder as expected from purely steric repulsion.^44^ Therefore, steric repulsion between the crowding in the cell and the H-NS constructs may well increase its oligomerization. In this scenario, crowders are depleted from the space between the H-NS dimers, providing a colloidal pressure difference that pushes the dimers toward higher oligomers. As the constructs have similar FRET values after the crowding increase, they probably have similar assembly structures, and the FRET increase is not due to DNA binding. While we do not have precise structural data, we can hypothesize that FRET increases by, for example, assembling linear oligomers to form fibers, or the disordered domain starting from amino acids 45, 65, or 83,^24,31,45^ generates fuzzy complexes leading to larger clusters. Crowding-triggered protein self-association in yeast and mammalian cells has been previously described for condensate-forming proteins,^10–12^ suggesting that cluster formation may be especially sensitive to crowding, which may happen here as well.

The relatively slow readjustment of H-NS oligomerization after Penicillin G addition could be a consequence of slow dissociation kinetics, semi-stable intermediate structures, or competitive binding with other molecules. Especially in the case of full-length H-NS, the DNA binding may reduce the re-assembly rate. While the mechanism is at this stage unclear, if we assume that these timeframes are more general to other protein-protein interactions in the cell, it would imply that the biochemical organization readjusts over hours after a sudden increase in macromolecular crowding. Macromolecular crowding relates to the biochemical organization: it can be expected that increased self-assembly of crowders reduces crowder volume and number density, reducing the crowding effect, which would, in turn, reduce crowder self-assembly, and so on until an equilibrium is reached. Therefore, one may extrapolate that protein self-assembly and crowding values both would equilibrate. Here, we indeed see that after the sudden increase, the macromolecular crowding slowly decreases,^35^ and the self-assembly increases to equilibrate over the time course of hours. We previously showed that adaptation to osmotic stress resulted in a slow decrease of crowding to reach equilibrium in the order of hours and was observed as an hours-long re-equilibration between the cell volume and crowding values.^46^ Therefore, the biochemical organization may respond slowly to stresses to find a new equilibrium with crowding.

Our observations on the H-NS sensitivity to macromolecular crowding are likely not exclusive to this protein. Other proteins of the bacterial histone family in *E. coli, S. typhimurium*, and *P. aeruginosa* have a large similarity in their architecture and may display crowding sensitivity.^27,28,33,34,47^ Next to DNA-binding proteins, other linear oligomers such as FtsZ increase their assembly by macromolecular crowding in buffer^44^ and, therefore, possibly also in cells. Crowding also enhances the self-assembly of proteins in mammalian cells, especially under more crowded osmotic stress conditions.^10,11^ Together, the data presented here show that crowding is likely involved in the self-organization of native proteins in *E. coli*, demonstrating the importance of crowding for cell physiology.

## Method details

### Growth conditions of *E. coli* cells

We used *E. coli* BL21 (DE3) for every experiment. The synthetic genes H-NS-VC, H-NS_64_-VC, and H-NS_86_-VC were codon optimized for *E. coli* and subcloned into the pBAD plasmid at the NcoI and HindIII restriction sites. Genes and plasmids, including VC control in the plasmid pRSET-A, were obtained from GeneArt. Bacteria were transformed with the VC constructs. A single colony of freshly transformed cells from an agar plate was grown overnight in MOPS minimal medium^48^ with 50 μg/mL ampicillin at 37 °C and 180 rpm. The cell cultures were then diluted to an OD_600_ of 0.03 in fresh MOPS minimal medium until they reached an OD_600_ = 0.1. Protein synthesis was induced by the addition of 5 mM arabinose and incubating for 3 h at 37 °C and 180 rpm. During protein synthesis, we maintained the cells in the exponential phase by keeping the OD_600_ between 0.1-0.3 by diluting with pre-warmed MOPS minimal medium.

### *In vitro* characterization of VC constructs

Cells containing either H-NS-VC or H-NS_86_-VC were harvested for 20 min at 4000 x *g* and 4 °C. The pellet was washed twice with isosmotic NaPi (100 mM, pH 7), weighed, and frozen at −80 °C overnight. The following day the cell pellet was thawed on ice, lysis buffer (30 mM K-Glutamate, 6 mM MgCl_2_, EDTA-free precursors, isosmotic NaPi (100 mM, pH 7) was added, and the sample was resuspended by vortexing. The cell lysis was achieved using TissueLyser LT, QIAGEN with three rounds of vigorous (50 s^−1^) oscillation, 10 min duration each, and in between, the sample was thawed on ice to avoid overheating. DNAseI (5 mg/mL) was added, and the sample was incubated for 25 min. The lysate was centrifuged (35 min, 21000 x *g*, 4 °C) until it was clear. The lysates containing the fluorescent constructs were then transferred to a MwCo 3kDA membrane and centrifuged (20 min, 4000 x *g*, 4 °C) to concentrate the proteins. About 15 μL of the concentrate was mounted on glass microscopy slides for the study of protein behavior. To evaluate the self-association of the VC constructs we added crowders (15% Ficoll-70) or salts (500 mM potassium glutamate or 50 mM magnesium chloride). The compounds were added in the same lysis buffer as the proteins.

### Preparation of *E. coli* lysozyme-spheroplasts

We created *E. coli* (± H-NS-VC, H-NS_64_-VC, H-NS_86_-VC) spheroplasts by modifying the protocol described in the literature.^49^ After 3 h of expression, 1 mL of the cell suspension was spun down at 3000 x *g* for 1 min. The pellet was resuspended in 500 μL 0.8 M sucrose solution. Then, we added the following solutions: 30 μL 1M Tris-HCl (pH 8.0), 24 μL 0.5 mg/mL lysozyme, (∼20 μg/mL final concentration), 6 μL 5 mg/mL DNase (∼50 μg/mL final concentration), and 6 μL 125 mM EDTA-NaOH (pH 8.0) (∼1.3 mM final concentration). Incubation of the sample for 10 min at 25 °C followed by 100 μL of STOP solution (10 mM Tris·HCl pH 8, 0.7 M sucrose, 20 mM MgCl_2_) was added to terminate the digestion. The cell suspension was then pelleted at 5000 x *rpm* for 5 min and resuspended in 50-70 μL of the spheroplasting mixture containing the STOP solution. 15 μL of the sample was mounted on glass microscopy slides for spheroplasts evaluation and quantification. For the adaptation of spheroplasts to lower osmolality, spheroplast suspensions were diluted to an osmolarity equal to MOPS medium (220 mOsmol/kg) by the gradual addition of Milli-Q water.

### Preparation of *E. coli* penicillin-spheroplasts

We created *E. coli* spheroplasts with Penicillin G based on literature protocols, with minor modifications.^50^ Briefly, 1 ml of cell suspension in the exponential growth phase was pelleted at 5000 x *rpm* for 5 min and resuspended in 10% w/v sucrose solution. Then we added the following solutions: 0.5 mg/mL sodium salt of Penicillin G solution and 0.2% w/v MgSO_4_. The sample was incubated for 1 h. Next, 15 μl of cell suspension was mounted on glass microscopy slides to evaluate and quantify spheroplasts.

### Microscopy settings and analysis

Before imaging, the cells were mixed with *E. coli* BL21 (DE3) cells without a plasmid, which were grown under the same conditions for background correction. For imaging bacteria, we mounted the glass microscopy slides on a Leica TCS SP8 laser-scanning confocal microscope. The mVenus in the VC constructs was excited using a 488-nm Argon laser, and the emission was split into a 505-555 nm and a 600-700 nm channel for the Donor (mVenus) and FRET (mCherry) channel, respectively.

Image analysis was performed with ImageJ, an open-source scientific image processing program.^51^ At first, intensity values of blank cells were subtracted from each, Donor and FRET channel. Subtracted values were then divided, and the FRET/Donor ratio of the VC constructs was calculated for single cells. The median value was estimated for each biological replicate, and then the average of three medians and the standard deviation was calculated.

### Calculation of H-NS_86_-VC foci size

We used the plot profile feature of ImageJ in the direction of the longitudinal axis of a single bacterium. The single wide peak that appeared corresponded to the H-NS_86_-VC foci and determined the diameter accordingly.

### Heat maps of VC constructs

To create heat maps of the fluorescent intensity of the constructs, we transferred the matrix with the absolute intensity values of each pixel of the FRET channel in Origin 2023. Heat maps were created, and background values were set as grey. We set the minimum fluorescence intensity to 3, as the average background from the cover slide is 2.5. HeatmapBio feature was selected to depict the distribution of protein densities through the entire cell. For the creation of the matrix of densities, we divided the maximum pixel value (in the intracellular space) with the minimum value, which corresponds to the external medium (on average 2.5, condition dependent). The resulting graph depicts dense areas in the cytoplasm for each construct and the distribution and differences of protein densities among constructs and conditions.

## Acknowledgments

The work was funded by the ERC Consolidator Grant (PArtCell; no. 864528) and the Netherlands Organization for Scientific Research (NWO Vidi; 723.015.002). The authors thank Rebecca Stephani for conducting part of the experimental work.

## Author contributions

P.T. and A.J.B. conceived the project. A.J.B. and P.T. designed the experiments, and T.P. performed the experiments and analyzed the data. T.P. and A.J.B. co-wrote the paper.

## Supplemental information

**Figure S1.**
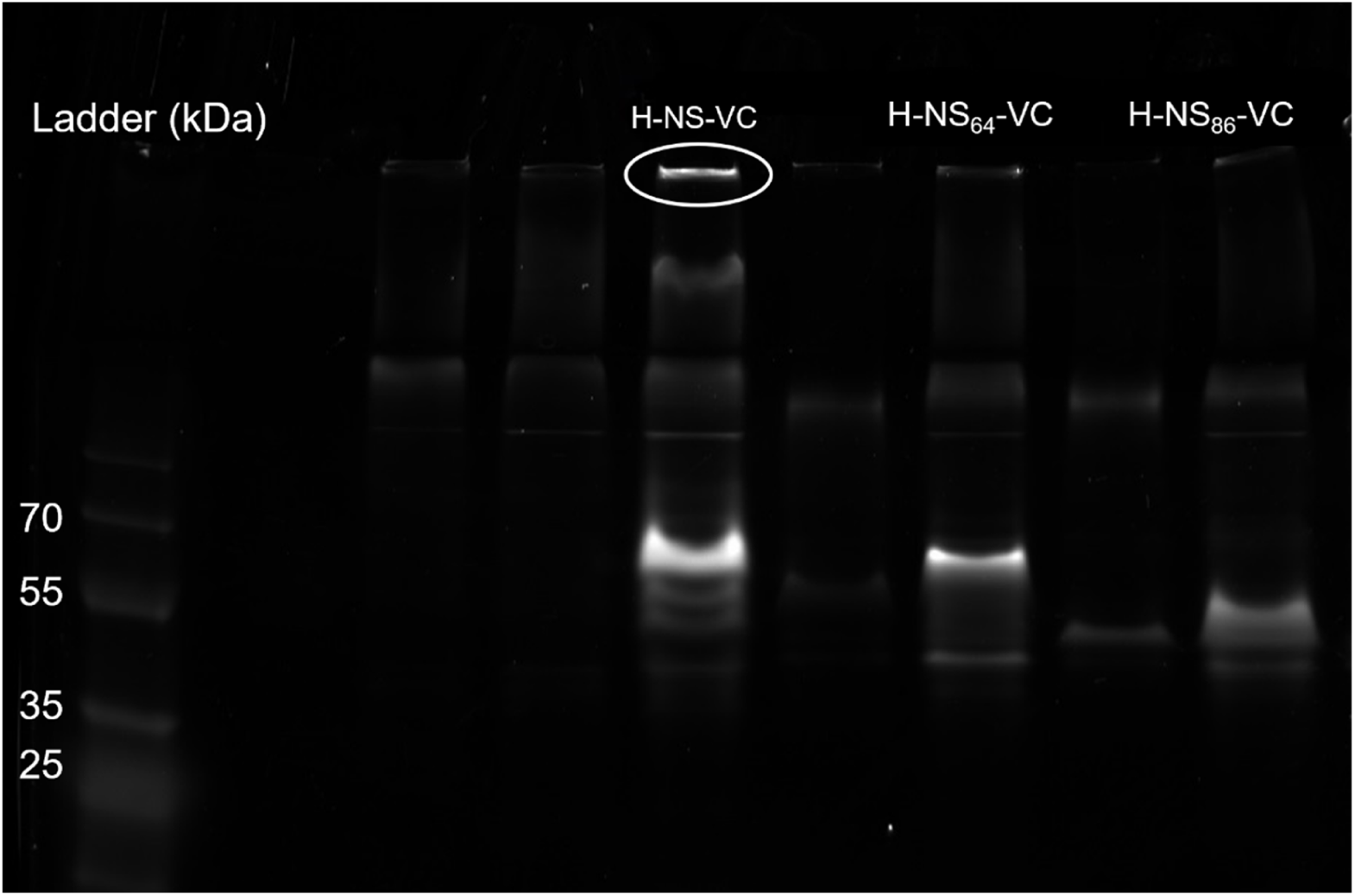
In-gel fluorescence (Alexa 647 laser) of the VC constructs. Native PAGE of unpurified H-NS-VC and its mutants in dilute cell lysates imaged by in gel-fluorescence with Alexa 647 imaging settings. All constructs show one major band corresponding to the protein and a few faint bands. Fluorescence in the wells for H-NS-VC is likely due to the formation of H-NS-VC-DNA complexes.

**Figure S2.**
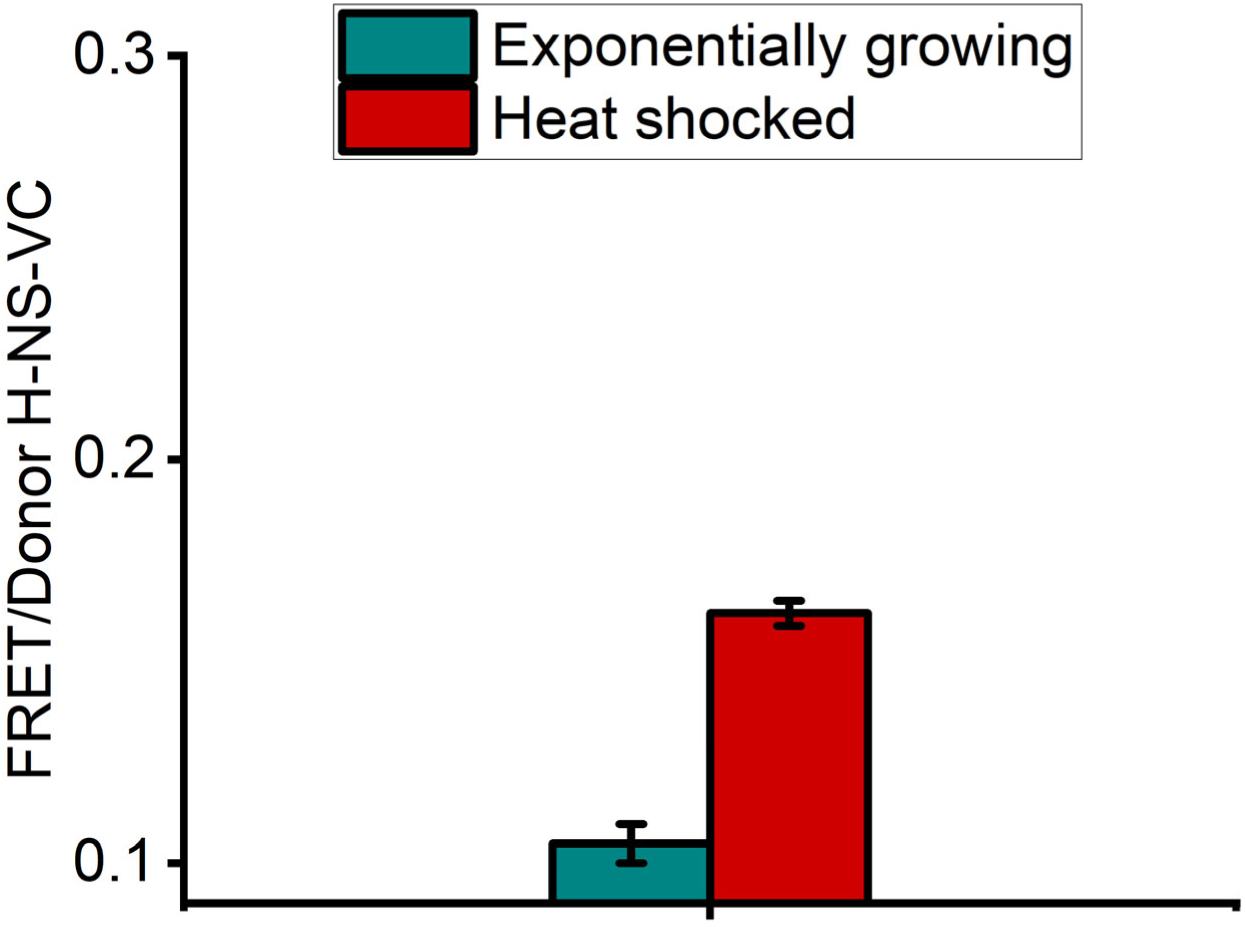
Heat shock increases H-NS-VC self-association. The FRET/Donor increases for H-NS-VC after heat shock at 42 °C for 6 min. Error bars represent the standard deviation of the average of three independent biological replicates.

**Figure S3.**
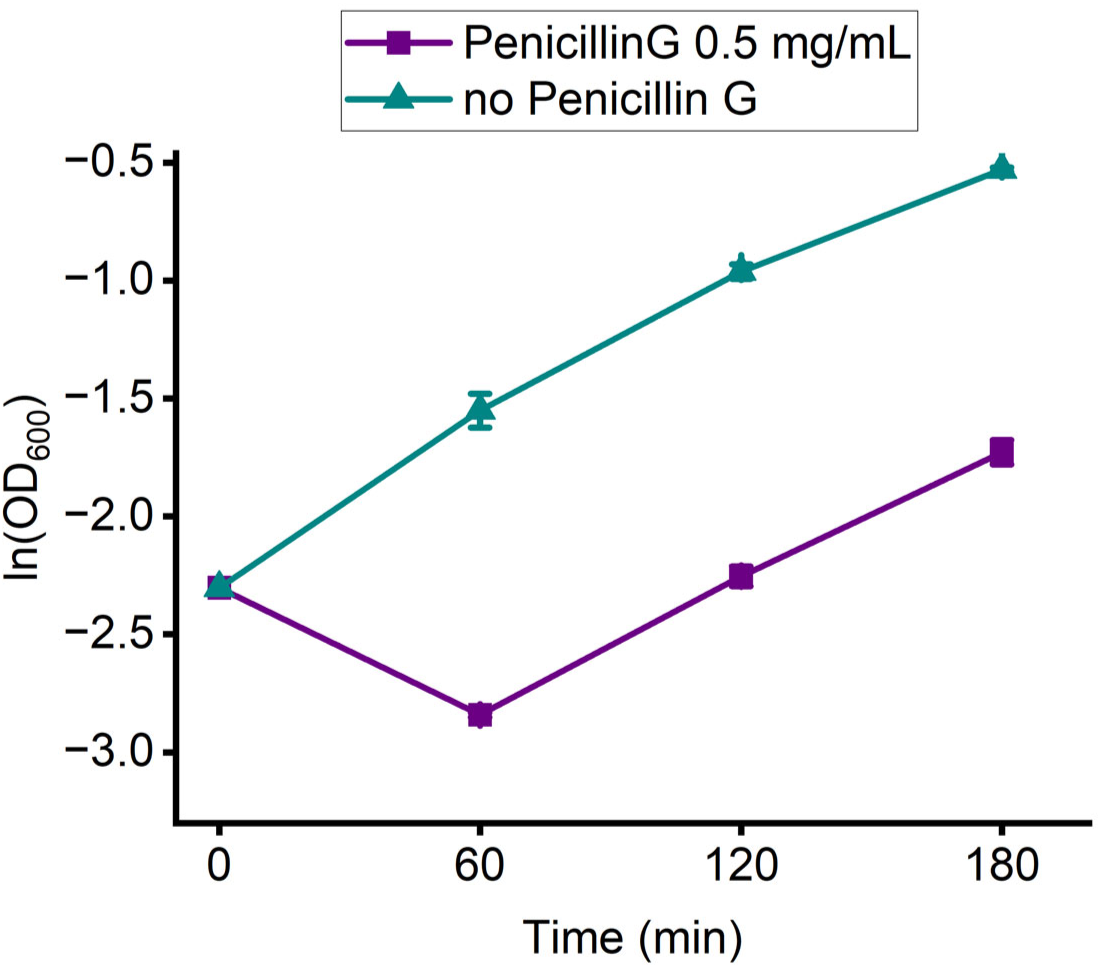
The growth curve of penicillin-treated cells. _lnOD600_ of exponentially growing cells in MOPS minimal medium and penicillin-treated cells. Error bars represent the standard deviation of the average of three independent biological replicates.

**Figure S4.**
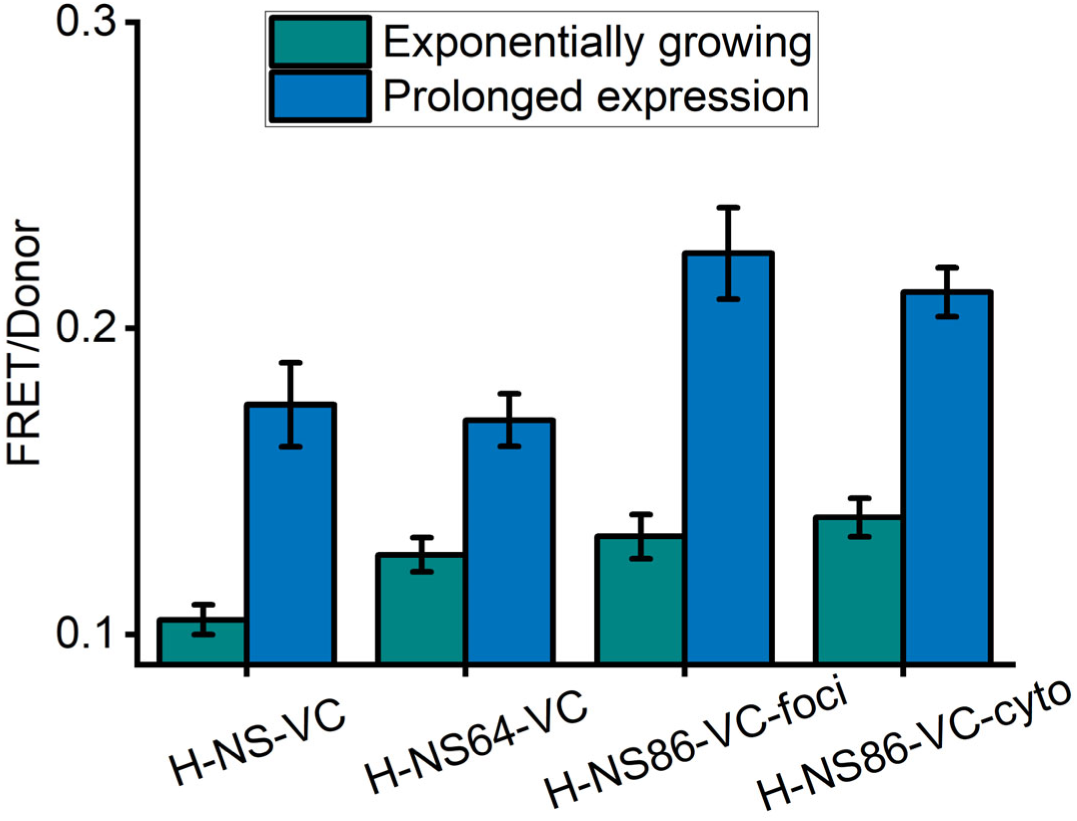
Self-association is increased with prolonged protein expression. The FRET/Donor ratio of VC constructs increases with prolonging protein expression from 3 to 6h. Error bars represent the standard deviation of the average of three independent biological replicates.

### DNA sequences of the constructs used in this study

#### VC (mVenus-mCherry)

ATGAAAGGTGAAGAGCTCTTCACTGGTGTTGTTCCAATTTTGGTTGAATTGGATGGTGATGTTAACGGCCATAAGTTTTCTGTTTCTGGTGAAGGTGAGGGTGATGCTACTTATGGTAAATTGACTTTGAAGTTGATCTGCACCACAGGTAAATTGCCAGTTCCCTGGCCAACTTTGGTTACTACTTTAGGTTACGGCTTGCAATGTTTTGCTAGATACCCAGATCATATGAAGCAACACGATTTCTTCAAATCCGCTATGCCAGAAGGTTACGTTCAAGAAAGAACTATCTTCTTCAAGGACGACGGTAACTACAAAACTAGAGCTGAAGTTAAGTTCGAAGGTGATACCTTGGTTAACAGGATTGAATTGAAGGGCATCGATTTCAAAGAGGATGGTAACATTTTGGGTCACAAGTTGGAGTACAACTACAACTCTCATAACGTTTACATTACCGCCGACAAGCAAAAGAATGGTATTAAGGCTAACTTCAAGATCAGGCACAACATTGAAGATGGTGGTGTTCAATTGGCTGATCACTATCAACAAAACACCCCAATTGGTGATGGTCCAGTTTTGTTGCCAGATAACCATTACTTGTCCTACCAGTCCAAGTTGTCTAAAGACCCAAACGAAAAAAGGGATCACATGGTTTTGTTGGAATTTGTTACAGCTGCTTCCGGTGATAACATGGCCATTATCAAAGAATTTATGAGGTTCAAGGTCCACATGGAAGGTTCTGTTAATGGTCACGAATTTGAGATTGAAGGTGAAGGCGAAGGTAGACCATATGAAGGTACTCAAACTGCTAAACTGAAGGTTACAAAAGGTGGTCCATTGCCATTTGCTTGGGATATTTTGTCTCCACAATTCATGTACGGTTCTAAGGCTTATGTAAAACACCCAGCTGATATCCCAGATTACTTGAAGTTGTCATTTCCAGAGGGTTTCAAGTGGGAAAGAGTTATGAATTTCGAGGATGGTGGCGTTGTTACTGTTACTCAAGATTCTTCATTACAGGACGGTGAGTTTATCTACAAGGTTAAGTTGAGAGGTACGAACTTTCCATCTGATGGTCCTGTTATGCAAAAAAAGACTATGGGTTGGGAAGCCTCTTCTGAAAGAATGTATCCAGAAGATGGCGCTTTGAAAGGTGAAATCAAACAAAGGTTGAAATTGAAGGACGGTGGTCATTATGATGCCGAAGTTAAGACTACTTACAAGGCTAAAAAGCCAGTTCAATTACCAGGTGCTTACAACGTCAACATCAAGTTGGATATCACTTCCCACAACGAAGATTACACCATCGTTGAACAATACGAAAGAGCTGAGGGTAGACATTCTACTGGTGGTATGTAA

#### H-NS-VC

ATGGGAAGCGAAGCGCTGAAAATTCTGAACAACATTCGCACCCTGCGCGCGCAGGCGCGCGAATGCACCCTGGAAACCCTGGAAGAAATGCTGGAAAAACTGGAAGTGGTGGTGAACGAACGCCGCGAAGAAGAAAGCGCGGCGGCGGCGGAAGTGGAAGAACGCACCCGCAAACTGCAGCAGTATCGCGAAATGCTGATTGCGGATGGCATTGATCCGAACGAACTGCTGAACAGCCTGGCGGCGGTGAAAAGCGGCACCAAAGCGAAACGCGCGCAGCGCCCGGCGAAATATAGCTATGTGGATGAAAACGGCGAAACCAAAACCTGGACCGGCCAGGGCCGCACCCCGGCGGTGATTAAAAAAGCGATGGATGAACAGGGCAAAAGCCTGGATGATTTTCTGATTAAACAGGGAGGTGGAGGCGGTTCCAAAGGTGAAGAGCTCTTCACTGGTGTTGTTCCAATTTTGGTTGAATTGGATGGTGATGTTAACGGCCATAAGTTTTCTGTTTCTGGTGAAGGTGAGGGTGATGCTACTTATGGTAAATTGACTTTGAAGTTGATCTGCACCACAGGTAAATTGCCAGTTcccTGGCCAACTTTGGTTACTACTTTAGGTTACGGCTTGCAATGTTTTGCTAGATACCCAGATCATATGAAGCAACACGATTTCTTCAAATCCGCTATGCCAGAAGGTTACGTTCAAGAAAGAACTATCTTCTTCAAGGACGACGGTAACTACAAAACTAGAGCTGAAGTTAAGTTCGAAGGTGATACCTTGGTTAACAGGATTGAATTGAAGGGCATCGATTTCAAAGAGGATGGTAACATTTTGGGTCACAAGTTGGAGTACAACTACAACTCTCATAACGTTTACATTACCGCCGACAAGCAAAAGAATGGTATTAAGGCTAACTTCAAGATCAGGCACAACATTGAAGATGGTGGTGTTCAATTGGCTGATCACTATCAACAAAACACCCCAATTGGTGATGGTCCAGTTTTGTTGCCAGATAACCATTACTTGTCCTACCAGTCCAAGTTGTCTAAAGACCCAAACGAAAAAAGGGATCACATGGTTTTGTTGGAATTTGTTACAGCTGCTTCCGGTGATAACATGGCCATTA TCAAAGAATTTATGAGGTTCAAGGTCCACATGGAAGGTTCTGTTAATGGTCACGAATTTGAGATTGAAGGTGAAGGCGAAGGTAGACCATATGAAGGTACTCAAACTGCTAAACTGAAGGTTACAAAAGGTGGTCCATTGCCATTTGCTTGGGATATTTTGTCTCCACAATTCATGTACGGTTCTAAGGCTTATGTAAAACACCCAGCTGATATCCCAGATTACTTGAAGTTGTCATTTCCAGAGGGTTTCAAGTGGGAAAGAGTTATGAATTTCGAGGATGGTGGCGTTGTTACTGTTACTCAAGATTCTTCATTACAGGACGGTGAGTTTATCTACAAGGTTAAGTTGAGAGGTACGAACTTTCCATCTGATGGTCCTGTTATGCAAAAAAAGACTATGGGTTGGGAAGCCTCTTCTGAAAGAATGTATCCAGAAGATGGCGCTTTGAAAGGTGAAATCAAACAAAGGTTGAAATTGAAGGACGGTGGTCATTATGATGCCGAAGTTAAGACTACTTACAAGGCTAAAAAGCCAGTTCAATTACCAGGTGCTTACAACGTCAACATCAAGTTGGATATCACTTCCCACAACGAAGATTACACCATCGTTGAACAATACGAAAGAGCTGAGGGTAGACATTCTACTGGTGGTATGTA A

#### H-NS_64_-VC

ATGGTGAGCGAAGCGCTGAAAATTCTGAACAACATTCGCACCCTGCGCGCGCAGGCGCGCGAATGCACCCTGGAAACCCTGGAAGAAATGCTGGAAAAACTGGAAGTGGTGGTGAACGAACGCCGCGAAGAAGAAAGCGCGGCGGCGGCGGAAGTGGAAGAACGCACCCGCAAACTGCAGCAGTATCGCGAAATGGGAGGTGGAGGCGGTTCCAAAGGTGAAGAGCTCTTCACTGGTGTTGTTCCAATTTTGGTTGAATTGGATGGTGATGTTAACGGCCATAAGTTTTCTGTTTCTGGTGAAGGTGAGGGTGATGCTACTTATGGTAAATTGACTTTGAAGTTGATCTGCACCACAGGTAAATTGCCAGTTcccTGGCCAACTTTGGTTACTACTTTAGGTTACGGCTTGCAATGTTTTGCTAGATACCCAGATCATATGAAGCAACACGATTTCTTCAAATCCGCTATGCCAGAAGGTTACGTTCAAGAAAGAACTATCTTCTTCAAGGACGACGGTAACTACAAAACTAGAGCTGAAGTTAAGTTCGAAGGTGATACCTTGGTTAACAGGATTGAATTGAAGGGCATCGATTTCAAAGAGGATGGTAACATTTTGGGTCACAAGTTGGAGTACAACTACAACTCTCATAACGTTTACATTACCGCCGACAAGCAAAAGAATGGTATTAAGGCTAACTTCAAGATCAGGCACAACATTGAAGATGGTGGTGTTCAATTGGCTGATCACTATCAACAAAACACCCCAATTGGTGATGGTCCAGTTTTGTTGCCAGATAACCATTACTTGTCCTACCAGTCCAAGTTGTCTAAAGACCCAAACGAAAAAAGGGATCACATGGTTTTGTTGGAATTTGTTACAGCTGCTTCCGGTGATAACATGGCCATTATCAAAGAATTTATGAGGTTCAAGGTCCACATGGAAGGTTCTGTTAATGGTCACGAATTTGAGATTGAAGGTGAAGGCGAAGGTAGACCATATGAAGGTACTCAAACTGCTAAACTGAAGGTTACAAAAGGTGGTCCATTGCCATTTGCTTGGGATATTTTGTCTCCACAATTCATGTACGGTTCTAAGGCTTATGTAAAACACCCAGCTGATATCCCAGATTACTTGAAGTTGTCATTTCCAGAGGGTTTCAAGTGGGAAAGAGTTATGAATTTCGAGGATGGTGGCGTTGTTACTGTTACTCAAGATTCTTCATTACAGGACGGTGAGTTTATCTACAAGGTTAAGTTGAGAGGTACGAACTTTCCATCTGATGGTCCTGTTATGCAAAAAAAGACTATGGGTTGGGAAGCCTCTTCTGAAAGAATGTATCCAGAAGATGGCGCTTTGAAAGGTGAAATCAAACAAAGGTTGAAATTGAAGGACGGTGGTCATTATGATGCCGAAGTTAAGACTACTTACAAGGCTAAAAAGCCAGTTCAATTACCAGGTGCTTACAACGTCAACATCAAGTTGGATATCACTTCCCACAACGAAGATTACACCATCGTTGAACAATACGAAAGAGCTGAGGGTAGACATTCTACTGGTGGTATGTAA

#### H-NS_86_-VC

ATGGTGAGCGAAGCGCTGAAAATTCTGAACAACATTCGCACCCTGCGCGCGCAGGCGCGCGAATGCACCCTGGAAACCCTGGAAGAAATGCTGGAAAAACTGGAAGTGGTGGTGAACGAACGCCGCGAAGAAGAAAGCGCGGCGGCGGCGGAAGTGGAAGAACGCACCCGCAAACTGCAGCAGTATCGCGAAATGCTGATTGCGGATGGCATTGATCCGAACGAACTGCTGAACAGCCTGGCGGCGGTGAAAAGCGGCACCGGAGGTGGAGGCGGTTCCAAAGGTGAAGAGCTCTTCACTGGTGTTGTTCCAATTTTGGTTGAATTGGATGGTGATGTTAACGGCCATAAGTTTTCTGTTTCTGGTGAAGGTGAGGGTGATGCTACTTATGGTAAATTGACTTTGAAGTTGATCTGCACCACAGGTAAATTGCCAGTTcccTGGCCAACTTTGGTTACTACTTTAGGTTACGGCTTGCAATGTTTTGCTAGATACCCAGATCATATGAAGCAACACGATTTCTTCAAATCCGCTATGCCAGAAGGTTACGTTCAAGAAAGAACTATCTTCTTCAAGGACGACGGTAACTACAAAACTAGAGCTGAAGTTAAGTTCGAAGGTGATACCTTGGTTAACAGGATTGAATTGAAGGGCATCGATTTCAAAGAGGATGGTAACATTTTGGGTCACAAGTTGGAGTACAACTACAACTCTCATAACGTTTACATTACCGCCGACAAGCAAAAGAATGGTATTAAGGCTAACTTCAAGATCAGGCACAACATTGAAGATGGTGGTGTTCAATTGGCTGATCACTATCAACAAAACACCCCAATTGGTGATGGTCCAGTTTTGTTGCCAGATAACCATTACTTGTCCTACCAGTCCAAGTTGTCTAAAGACCCAAACGAAAAAAGGGATCACATGGTTTTGTTGGAATTTGTTACAGCTGCTTCCGGTGATAACATGGCCATTATCAAAGAATTTATGAGGTTCAAGGTCCACATGGAAGGTTCTGTTAATGGTCACGAATTTGAGATTGAAGGTGAAGGCGAAGGTAGACCATATGAAGGTACTCAAACTGCTAAACTGAAGGTTACAAAAGGTGGTCCATTGCCATTTGCTTGGGATATTTTGTCTCCACAATTCATGTACGGTTCTAAGGCTTATGTAAAACACCCAGCTGATATCCCAGATTACTTGAAGTTGTCATTTCCAGAGGGTTTCAAGTGGGAAAGAGTTATGAATTTCGAGGATGGTGGCGTTGTTACTGTTACTCAAGATTCTTCATTACAGGACGGTGAGTTTATCTACAAGGTTAAGTTGAGAGGTACGAACTTTCCATCTGATGGTCCTGTTATGCAAAAAAAGACTATGGGTTGGGAAGCCTCTTCTGAAAGAATGTATCCAGAAGATGGCGCTTTGAAAGGTGAAATCAAACAAAGGTTGAAATTGAAGGACGGTGGTCATTATGATGCCGAAGTTAAGACTACTTACAAGGCTAAAAAGCCAGTTCAATTACCAGGTGCTTACAACGTCAACATCAAGTTGGATATCACTTCCCACAACGAAGATTACACCATCGTTGAACAATACGAAAGAGCTGAGGGTAGACATTCTACTGGTGGTATGTAA

#### crGE2.3 (mEGFP/mScarlet-I)

ATGAAAGGTGAAGAACTGTTTACCGGTGTTGTTCCGATTCTGGTTGAACTGGATGGTGACGTTAATGGTCACAAATTTTCAGTTAGCGGTGAAGGCGAAGGTGATGCAACCTATGGTAAACTGACCCTGAAATTTATCTGTACCACCGGCAAACTGCCGGTTCCGTGGCCGACACTGGTTACCACACTGACCTATGGTGTTCAGTGTTTTAGCCGTTATCCTGATCACATGAAACAGCACGATTTTTTCAAAAGCGCAATGCCGGAAGGTTATGTTCAAGAACGTACCATCTTCTTCAAAGATGACGGCAACTATAAAACCCGTGCCGAAGTTAAATTTGAAGGTGATACCCTGGTGAATCGCATTGAACTGAAAGGCATCGATTTTAAAGAGGATGGTAATATCCTGGGCCACAAACTGGAATATAATTATAATAGCCACAACGTGTACATCATGGCCGACAAACAGAAAAATGGCATCAAAGTGAACTTCAAGATCCGCCATAATATTGAAGATGGTTCAGTTCAGCTGGCCGATCATTATCAGCAGAATACCCCGATTGGTGATGGTCCGGTTCTGCTGCCGGATAATCATTATCTGAGCACCCAGAGCAAACTGAGCAAAGATCCGAATGAAAAACGCGATCACATGGTGCTGCTGGAATTTGTTACCGCAGCAGGTATTACCTTAGGTATGGATGAACTGTATAAAGGATCCGGTGGTAGCGGTGGTTCAGGTGGTAGTGGCGGTAGTGGTGGCAGCGGTGCAGAAGCAGCAGCAAAAGAAGCCGCTGCCAAAGAAGCGGCAGCGAAAGAGGCTGCCGCAAAAGAGGCAGCAGCGAAAGAAGCAGCGGCTAAAGCAGGTTCAGGCGGAAGCGGAGGCAGTGGTGGATCAGGCGGATCTGGTGGCTCAGGTGCCGAGGCAGCAGCAAAAGAGGCAGCTGCTAAAG AGGCTGCTGCAAAAGAAGCAGCCGCAAAAGAGGCAGCGGCAAAAGAAGCCGCAGCAAAAGCAGGTAGTGGTGGAAGTGGCGGTTCCGGTGGCTCTGGTGGAAGCGGTGGCTCCGGAGAGCTCGTTAGTAAAGGCGAAGCAGTTATTAAAGAATTTATGCGCTTCAAAGTGCACATGGAAGGTAGCATGAATGGCCATGAATTTGAAATCGAAGGTGAAGGTGAGGGTCGTCCGTATGAAGGCACCCAGACCGCAAAACTGAAAGTTACCAAAGGTGGTCCGCTGCCGTTTAGCTGGGATATTCTGAGTCCGCAGTTTATGTATGGTAGCCGTGCATTTATCAAACATCCGGCAGATATCCCGGATTATTACAAACAGAGCTTTCCCGAAGGTTTTAAATGGGAACGTGTGATGAATTTTGAGGATGGTGGTGCAGTTACCGTTACACAGGATACCAGCCTGGAAGATGGCACC CTGATCTATAAAGTTAAACTGCGTGGCACCAATTTTCCGCCAGATGGTCCTGTTATGCAG AAAAAAACCATGGGTTGGGAAGCAAGCACCGAACGTCTGTATCCTGAAGATGGCGTTCT GAAAGGTGATATCAAAATGGCACTGCGTCTGAAAGATGGTGGTCGTTATCTGGCAGATTT CAAAACCACCTACAAAGCCAAAAAACCGGTTCAGATGCCTGGTGCATATAATGTTGATC GCAAACTGGATATCACCAGCCATAATGAAGATTATACCGTGGTGGAACAGTATGAACGT AGCGAAGGTCGTCATAGTACCGGTGGCATGGATGAATTATACAAAGGTGGCACCTAA

## References

1. Stefanis, L. α-Synuclein in Parkinson’s Disease. Cold Spring Harb. Perspect. Med. 2, a009399 (2012).

2. Zimmerman, S. B. & Trach, S. O. Estimation of macromolecule concentrations and excluded volume effects for the cytoplasm of Escherichia coli. J. Mol. Biol. 222, 599–620 (1991).

3. Cayley, S., Lewis, B. A., Guttman, H. J. & Record, M. T. Characterization of the cytoplasm of Escherichia coli K-12 as a function of external osmolarity: Implications for protein-DNA interactions in vivo. J. Mol. Biol. 222, 281–300 (1991).

4. Rivas, G. & Minton, A. P. Influence of Nonspecific Interactions on Protein Associations: Implications for Biochemistry In Vivo. Annu. Rev. Biochem. 91, 321–351 (2022).

5. Yamin, G. et al. Forcing Nonamyloidogenic β-Synuclein To Fibrillate. Biochemistry 44, 9096–9107 (2005).

6. Munishkina, L. A., Cooper, E. M., Uversky, V. N. & Fink, A. L. The effect of macromolecular crowding on protein aggregation and amyloid fibril formation. J. Mol. Recognit. 17, 456–464 (2004).

7. Alric, B., Formosa-Dague, C., Dague, E., Holt, L. J. & Delarue, M. Macromolecular crowding limits growth under pressure. Nat. Phys. 18, 411–416 (2022).

8. Boersma, A. J., Zuhorn, I. S. & Poolman, B. A sensor for quantification of macromolecular crowding in living cells. Nat. Methods 12, 227–229 (2015).

9. Joyner, R. P. et al. A glucose-starvation response regulates the diffusion of macromolecules. eLife 5, e09376 (2016).

10. Watanabe, K. et al. Cells recognize osmotic stress through liquid–liquid phase separation lubricated with poly(ADP-ribose). Nat. Commun. 12, 1353 (2021).

11. Delarue, M. et al. mTORC1 Controls Phase Separation and the Biophysical Properties of the Cytoplasm by Tuning Crowding. Cell 174, 338-349.e20 (2018).

12. Boyd-Shiwarski, C. R. et al. WNK kinases sense molecular crowding and rescue cell volume via phase separation. Cell 185, 4488-4506.e20 (2022).

13. Wei, S.-P. et al. Formation and functionalization of membraneless compartments in Escherichia coli. Nat. Chem. Biol. 16, 1143–1148 (2020).

14. Sang, D. et al. Condensed-phase signaling can expand kinase specificity and respond to macromolecular crowding. Mol. Cell 82, 3693-3711.e10 (2022).

15. Yeong, V., Werth, E. G., Brown, L. M. & Obermeyer, A. C. Formation of Biomolecular Condensates in Bacteria by Tuning Protein Electrostatics. ACS Cent. Sci. 6, 2301–2310 (2020).

16. Varshavsky, A. J., Nedospasov, S. A., Bakayev, V. V., Bakayeva, T. G. & Georgiev, G. P. Histone-like proteins in the purified Escherichia coli deoxyribonucleoprotein. Nucleic Acids Res. 4, 2725–2746 (1977).

17. Tupper, A. e. et al. The chromatin-associated protein H-NS alters DNA topology in vitro. EMBO J. 13, 258–268 (1994).

18. Dame, R. T. The role of nucleoid-associated proteins in the organization and compaction of bacterial chromatin. Mol. Microbiol. 56, 858–870 (2005).

19. van der Valk, R. A. et al. Mechanism of environmentally driven conformational changes that modulate H-NS DNA-bridging activity. eLife 6, e27369 (2017).

20. Liu, Y., Chen, H., Kenney, L. J. & Yan, J. A divalent switch drives H-NS/DNA-binding conformations between stiffening and bridging modes. Genes Dev. 24, 339–344 (2010).

21. Dame, R. T., Wyman, C. & Goosen, N. H-NS mediated compaction of DNA visualised by atomic force microscopy. Nucleic Acids Res. 28, 3504–3510 (2000).

22. Shindo, H. et al. Solution structure of the DNA binding domain of a nucleoid-associated protein, H-NS, from Escherichia coli. FEBS Lett. 360, 125–131 (1995).

23. Dame, R. T., Noom, M. C. & Wuite, G. J. L. Bacterial chromatin organization by H-NS protein unravelled using dual DNA manipulation. Nature 444, 387–390 (2006).

24. Bloch, V. et al. The H-NS dimerization domain defines a new fold contributing to DNA recognition. Nat. Struct. Mol. Biol. 10, 212–218 (2003).

25. Grainger, D. C. Structure and function of bacterial H-NS protein. Biochem. Soc. Trans. 44, 1561–1569 (2016).

26. Winardhi, R. S., Yan, J. & Kenney, L. J. H-NS Regulates Gene Expression and Compacts the Nucleoid: Insights from Single-Molecule Experiments. Biophys. J. 109, 1321–1329 (2015).

27. Smyth, C. P. et al. Oligomerization of the chromatin-structuring protein H-NS. Mol. Microbiol. 36, 962–972 (2000).

28. Rimsky, S. Structure of the histone-like protein H-NS and its role in regulation and genome superstructure. Curr. Opin. Microbiol. 7, 109–114 (2004).

29. Dorman, C. J. H-NS: a universal regulator for a dynamic genome. Nat. Rev. Microbiol. 2, 391–400 (2004).

30. Ueguchi, C., Seto, C., Suzuki, T. & Mizuno, T. Clarification of the dimerization domain and its functional significance for the Escherichia coli nucleoid protein H-NS 11Edited by I. B. Holland. J. Mol. Biol. 274, 145–151 (1997).

31. Gao, Y. et al. Charged residues in the H-NS linker drive DNA binding and gene silencing in single cells. Proc. Natl. Acad. Sci. 114, 12560–12565 (2017).

32. Esposito, D. et al. H-NS Oligomerization Domain Structure Reveals the Mechanism for High Order Self-association of the Intact Protein. J. Mol. Biol. 324, 841–850 (2002).

33. Arold, S. T., Leonard, P. G., Parkinson, G. N. & Ladbury, J. E. H-NS forms a superhelical protein scaffold for DNA condensation. Proc. Natl. Acad. Sci. 107, 15728–15732 (2010).

34. Qin, L. et al. Structural basis for osmotic regulation of the DNA binding properties of H-NS proteins. Nucleic Acids Res. 48, 2156–2172 (2020).

35. Pittas, T., Zuo, W. & Boersma, A. J. Cell wall damage increases macromolecular crowding effects in the Escherichia coli cytoplasm. 2022.10.03.510584 Preprint at https://doi.org/10.1101/2022.10.03.510584 (2022).

36. Wan, Q., Mouton, S. N., Veenhoff, L. M. & Boersma, A. J. A FRET-based method for monitoring structural transitions in protein self-organization. Cell Rep. Methods 2, 100184 (2022).

37. Ma, Y. et al. An intermolecular FRET sensor detects the dynamics of T cell receptor clustering. Nat. Commun. 8, 15100 (2017).

38. Büning, S. et al. Conformational dynamics and self-association of intrinsically disordered Huntingtin exon 1 in cells. Phys. Chem. Chem. Phys. 19, 10738–10747 (2017).

39. Caron, N. S., Desmond, C. R., Xia, J. & Truant, R. Polyglutamine domain flexibility mediates the proximity between flanking sequences in huntingtin. Proc. Natl. Acad. Sci. 110, 14610–14615 (2013).

40. Liu, Y. et al. A model for chromosome organization during the cell cycle in live E. coli. Sci. Rep. 5, 17133 (2015).

41. Ceschini, S. et al. Multimeric Self-assembly Equilibria Involving the Histone-like Protein H-NS: A THERMODYNAMIC STUDY*. J. Biol. Chem. 275, 729–734 (2000).

42. Rafiei, N., Cordova, M., Navarre, W. W. & Milstein, J. N. Growth Phase-Dependent Chromosome Condensation and Heat-Stable Nucleoid-Structuring Protein Redistribution in Escherichia coli under Osmotic Stress. J. Bacteriol. 201, e00469–19 (2019).

43. Will, W. R., Whitham, P. J., Reid, P. J. & Fang, F. C. Modulation of H-NS transcriptional silencing by magnesium. Nucleic Acids Res. 46, 5717–5725 (2018).

44. Rivas, G., Fernández, J. A. & Minton, A. P. Direct observation of the enhancement of noncooperative protein self-assembly by macromolecular crowding: Indefinite linear self-association of bacterial cell division protein FtsZ. Proc. Natl. Acad. Sci. 98, 3150–3155 (2001).

45. Zhao, X. et al. Molecular basis for the adaptive evolution of environment-sensing by H-NS proteins. eLife 10, e57467.

46. Liu, B., Hasrat, Z., Poolman, B. & Boersma, A. J. Decreased Effective Macromolecular Crowding in Escherichia coli Adapted to Hyperosmotic Stress. J. Bacteriol. 201, e00708–18 (2019).

47. Suzuki, C. et al. Oligomerization Mechanisms of an H-NS Family Protein, Pmr, Encoded on the Plasmid pCAR1 Provide a Molecular Basis for Functions of H-NS Family Members. PLOS ONE 9, e105656 (2014).

48. Neidhardt Frederick C., Bloch Philip L., & Smith David F. Culture Medium for Enterobacteria. J. Bacteriol. 119, 736–747 (1974).

49. Renner, L. D. & Weibel, D. B. Cardiolipin microdomains localize to negatively curved regions of Escherichia coli membranes. Proc. Natl. Acad. Sci. 108, 6264–6269 (2011).

50. Hirokawa, H. Biochemical and cytological observations during the reversing process from spheroplasts to rod-form cells in escherichia coli. J. Bacteriol. 84, 1161–1168 (1962).

51. Schneider, C. A., Rasband, W. S. & Eliceiri, K. W. NIH Image to ImageJ: 25 years of image analysis. Nat. Methods 9, 671–675 (2012).

